# Entecavir resistance mutations rtL180M/T184L/M204V combined with rtA200V lead to tenofovir resistance

**DOI:** 10.1101/632984

**Authors:** Dong Jiang, Jianghua Wang, Xuesen Zhao, Yuxin Li, Qun Zhang, Chuan Song, Hui Zeng, Xianbo Wang

## Abstract

**Background & Aims:** Tenofovir disoproxil fumarate (TDF) imposes a high genetic barrier to drug resistance and potently inhibits replication of multidrug-resistant hepatitis B virus (MDR HBV) and few clinical cases with confirmed TDF-resistance have been reported to date. We reported a quadruple mutant which showed moderate resistant to TDF.

**Methods and results:** Viral rebound was reported in a patient with chronic hepatitis B who underwent TDF monotherapy and harbored a quadruple mutant consisting of classic ETV-resistance mutations (rtL180M/T184L/M204V) together with an rtA200V mutation in the reverse transcriptase gene. Sequencing analysis revealed that this quadruple mutant emerged as a major viral population. *In vitro* phenotyping demonstrated that the rtL180M/T184L/A200V/M204V mutant had moderate resistance to TDF treatment, with a 4.52-fold higher half maximal effective concentration than that of wild-type virus. Importantly, this patient with TDF resistance achieved virological suppression after TDF/ETV combination rescue therapy.

**Conclusion:** An rtL180M/T184L/A200V/M204V mutant with moderate resistance to TDF monotherapy was selected during sequential NA treatment in a stepwise manner. ETV/TDF combination therapy effectively suppressed replication of this TDF resistant mutant. Our studies provide novel insights into the treatment of NA-naïve patients as well as patients with TDF resistance.

## Introduction

Nearly 350 million individuals are chronically infected with hepatitis B virus (HBV) worldwide. Approximately 2% to 4% of untreated patients with HBeAg-positive or HBeAg-negative hepatitis develop cirrhosis annually, and one-third of these patients eventually progress to fatal liver diseases.(1) Since lamivudine (LAM) was approved by the FDA in 1998, NAs have been widely used to treat chronic hepatitis B (CHB) and markedly improve the clinical outcome of this patient population. However, drug resistance to NAs has emerged as a critical issue during long-term therapy.(2) It has been well documented that LAM resistance occur at a frequency as high as approximately 20% per year.(3) Compared with LAM, adefovir dipivoxil (ADV) has a relatively lower resistance rate (approximately 30% at the end of a 5-year treatment).(4) Of note, entecavir (ETV), imposes a higher genetic barrier to drug resistance.(5–7) ETV resistance occured in only 1.2% of NA naïve patients, whereas the frequency is increased to 51% in patients infected with LAM-resistant strains during a 5-year treatment period.(7)

More recently, tenofovir disoproxil fumarate (TDF) has been shown to be potently effective against different HBV genotypes as well as LAM-, ADV- or ETV-resistant mutants.(8–11) Kitrinos and Liu have reported an absence of TDF resistance in patients even after 6 years of TDF monotherapy.(12) Although sporadic cases with viral breakthrough during TDF treatment have been reported by several studies, most of these findings resulted from non-compliance with the anti-viral treatment.(10) Recently, Shirvani-Dastgerdi has identified an uncommon pattern of rtS78T/sC69* mutation in two TDF patients, which was associated with a 1.6-fold change in the half maximal effective concentration (EC50).(13) However, the presence of TDF resistant mutants together with clinical viral breakthrough still turn into a rare event.

In the present study, we identified a quadruple HBV mutant (rtL180M/T184L/A200V/M204V) in a CHB patient with clinical viral rebound during TDF monotherapy. *In vitro* study indicated that this quadruple HBV mutant exhibited decreased sensitivity to TDF, with an approximate 4.5-fold higher EC50 than that of wild-type virus. Importantly this mutant was virologically suppressed by subsequent TDF/ETV combination rescue therapy.

## Patient and Methods

### Patient

A 47-year-old woman who was diagnosed with CHB in 2001 and underwent NA treatment and serum levels of HBV DNA and ALT were periodically monitored. HBV DNA levels were monitored centrally with a Roche COBAS TaqMan HBV test(lower limit of quantification = 20 IU/mL). The HBV reverse transcriptase (RT) sequence was analyzed to identify potential drug resistance sites when virological breakthrough occurred. This study was approved by the Committee of Ethics at Beijing Ditan Hospital, Capital Medical University, and was conducted in accordance with the principles of the Declaration of Helsinki. All blood samples were collected after written informed consent was obtained.

### Extraction and amplification of HBV RT, direct sequencing and clonal analysis

HBV DNA was extracted from 50 μl of serum with a PureLink Viral RNA/DNA Mini Kit (Invitrogen) and collected in 20 μl of water. HBV DNA amplification was done with Platinum SuperFi DNA Polymerase (Invitrogen) with primers P1 (CTCTCCACCTCTAAGAGACA, 3177-3196) and P8 (CTCCAGACCGGCTGCGAGCA, 1318-1297). PCR products were purified with a Wizard SV Gel and PCR Clean-Up System (Promega) and directly sequenced with an Applied Biosystems 3730XL instrument. HBV quasispecies were assessed by clonal analysis. The PCR fragment of HBV RT was cloned into the pGEM-T easy vector (Promega). Clones bearing the insert were sequenced and analyzed on the basis of alignment with the HBV wild type sequence.

### Construction of HBV plasmids harboring rtL180M/A200V/T184L/M204V mutations

Single or multi-site mutations of rtL180M/A200V/T184L/M204V were introduced by PCR based site-directed mutagenesis into pUC18-HBV1.2-WT carrying a 1.2-fold full-length genotype C HBV genome.(14) Plasmids bearing different sequencing-confirmed mutations were extracted with a NucleoBond Xtra Midi Plus plasmid purification kit (MACHEREY-NAGEL GmbH & Co. KG).

### Compounds

ETV and TDF compounds were provided by Sunshine Lake Pharma, China. Their purities were ≥98%, on the basis of HPLC analysis. Compounds were dissolved in DMSO to generate 10 mM stock solutions, which were stored at −80°C until use.

### Cell culture and transient transfection of HBV

For transfection, six-well culture plates were seeded with 8×10^5^ HepG2 cells per well in DMEM/F12 (Thermo fisher) supplemented with 10% (vol/vol) fetal bovine serum. Sixteen hours post-seeding, cells were transfected with 2 μg of plasmid DNA with Fugene HD transfection reagent (Promega). Six hours later, cell culture medium was added with TDF (final concentrations 0, 1, 2, 4, 8, 16, 32 and 64 nM). Culture supernatants and cells were harvested after 96 h of treatment. Transfection efficiency was measured through co-transfection of the reporter plasmid pmirGLO (Promega). Luciferase activity was measured with a Bright-Glo Luciferase Assay System (Promega) after the cells were lysed with lysis buffer. Transfection experiments were performed three times.

### DNA extraction and Southern blotting of viral DNA

Four hundred microliters of Core DNA lysis buffer (10 mM Tris-HCl, pH 8.0, 1 mM EDTA and 1% NP-40) was added to the cells. The cells were rocked on ice for 20 minutes, and nuclei and debris were removed by centrifugation at 10,000 rpm for 5 minutes at 4°C. The supernatant was treated with 2 U of Turbo DNase I (Thermo fisher) for 1 h at 37°C. Then the DNase was inactivated by 15 mM of EDTA and heating for 10 minutes at 75°C. Cell lysate was digested with 200 μg/mL proteinase K at 55°C for 1 h, then extracted twice with phenol/chloroform/isoamyl alcohol. HBV DNA was precipitated with a 1/10 volume of 3 M NaOAc and an equal volume of isopropanol at −20°C overnight. Pellets containing HBV replication intermediate DNA were suspended in 20 μl water. The intracellular HBV DNA was detected by Southern blotting. Blotted membranes were hybridized and detected with DIG Northern Starter Kit(Roche). Chemiluminescence signals were detected with a ChemiDoc MP Imaging System (Bio-Rad) and analyzed in Image Lab software Version 5.2.1 (Bio-Rad).

### Homology modeling of the HBV RT-DNA complex

A homology model of the HBV polymerase-DNA-tenofovir triphosphate complex was constructed in PyMOL software, on the basis of the sequence alignment between HBV polymerase (GenBank accession number: AAR99338.1) and HIV-1 RT, and the crystal structure of the HIV-1 RT-DNA-dNTP complex (Protein Data Bank accession number 1RTD). Coot® software was used to model the rtL180M/T184L/A200V/M204V mutations in HBV RT.

### Statistical analysis

HBV replication intermediates were analyzed and quantified in Image Lab software. Results are expressed as mean±standard deviation. EC50 was calculated through non-linear regression analysis in GraphPad Prism version 5.01 for Windows (GraphPad Software, San Diego, CA, USA).

## Results

### Procedure and duration of treatment with LAM, ADV, ETV and TDF, alone or in combination

A 47-year-old woman was diagnosed with CHB and initially received standard LAM treatment (100 mg QD) in 2001. Three years later, ADV (10 mg QD) was added to the LAM treatment regimen. In March 2011, a LAM resistant mutant rtM204V was detected in HBV DNA from the patient’s serum; thus, the LAM/ADV treatment was switched to ETV (1.0 mg QD) plus ADV (10 mg QD). The level of ALT declined to below normal limit, and the viral load was between 8×10^3^ and 1×10^4^ IU/ml (Fig. 1).

**Fig. 1.**
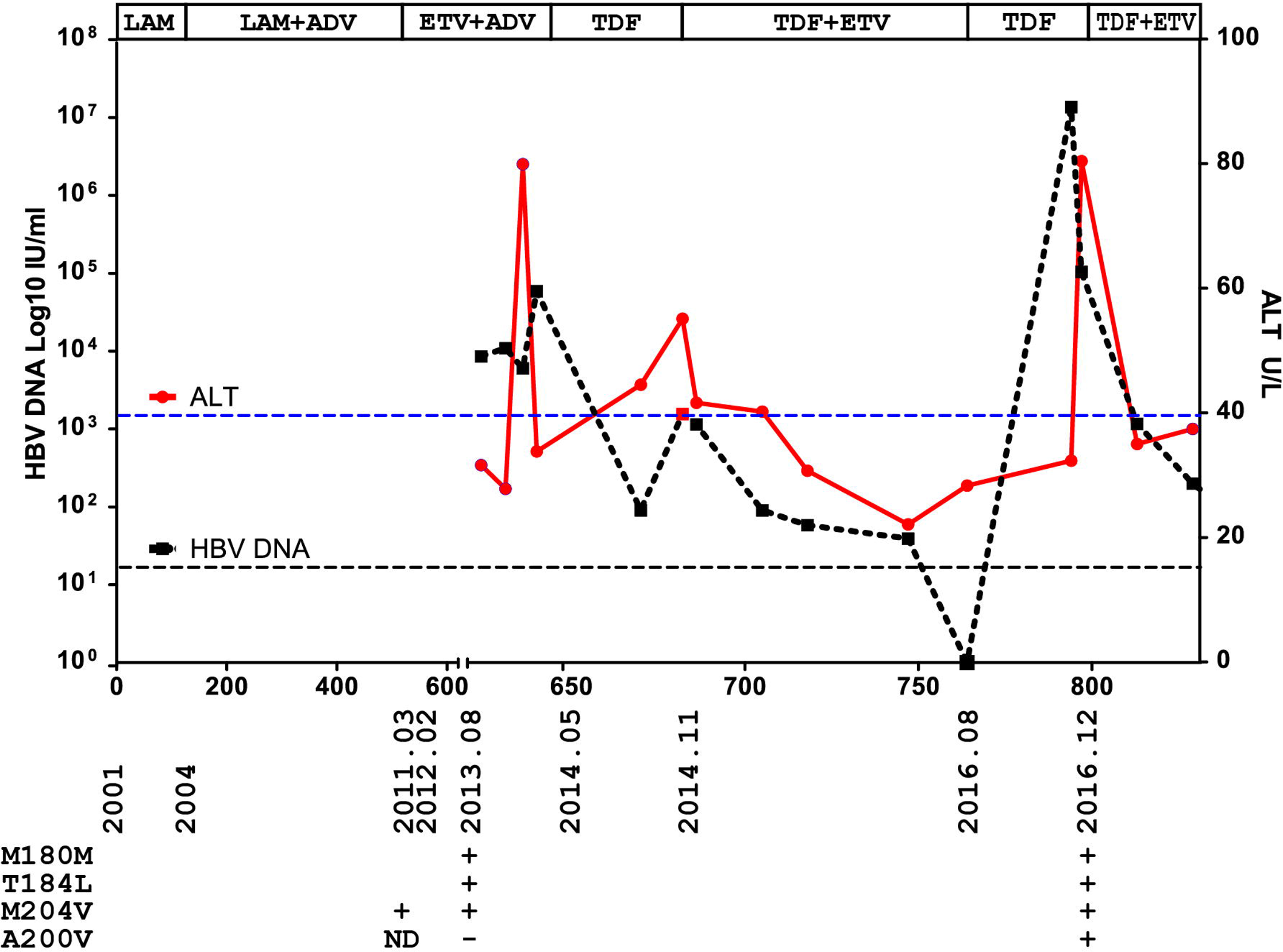
Patient clinical course and mutation information. HBV DNA viral load and ALT levels in relation to NA treatment are shown. The HBV DNA detection limit and normal ALT level are indicated as dashed black and red lines, respectively. The treatment or sampling time points for mutation analyses and corresponding mutation patterns are indicated.

In 2013, given the increase in serum ALT level to 80 U/L, a sequencing analysis was performed, and an ETV-resistant triple-mutation of rtL180M/T184L/M204V was detected. In May 2014, the patient received TDF monotherapy (300 mg QD). Because of abnormal ALT level and detectable serum HBV DNA, TDF+ETV combination therapy was started in November 2014. During the subsequent 9 months’ treatment, the viral load gradually decreased and eventually became undetectable (<20 IU/ml), and ALT was normalized.

In August 2016, ETV was tentatively discontinued, and TDF monotherapy continued. Four months later, virological breakthrough occurred with an increased viral load (HBV DNA 10^7^ IU/ml) and elevated ALT (80 U/L). An ETV/TDF regimen was resumed. After 8 months’ treatment, the HBV viral load decreased to 1.16×10^3^ IU/ml in Aug. 2017, and the ALT level returned to normal (Fig. 1).

The patient was diagnosed with decompensate cirrhosis and experienced rupture of esophageal and gastric varices in 2010 and 2015. Splenomegaly was confirmed in July 2010, and splenectomy was performed in 2012. Ascites was found in August 2015. HBeAg seroconversion was confirmed in July 2012. The HBeAg became positive in August 2013 and has remained positive to date.

### The quadruple mutant rtL180M/T184L/A200V/M204V emerged as a major viral population

We assessed whether the viral breakthrough in this case was accompanied with potential TDF resistance-associated mutations. Sequencing analysis revealed classic ETF-resistance mutations (rtL180M/T184L/M204V) in December 2016. In addition, an rtA200V was detected (Fig. 2A). rtA200V was found only in four sequences among 8038 full-length HBV sequences in GenBank, indicating that rtA200V is a drug associated mutation rather than a polymorphic change. Other multi-drug resistance (MDR) mutations, including rtI169T, rtV173L, rtA181V/T, rtS202G/I, rtN236T and rtM250V, were not detected (Fig. 2B).

**Fig. 2.**
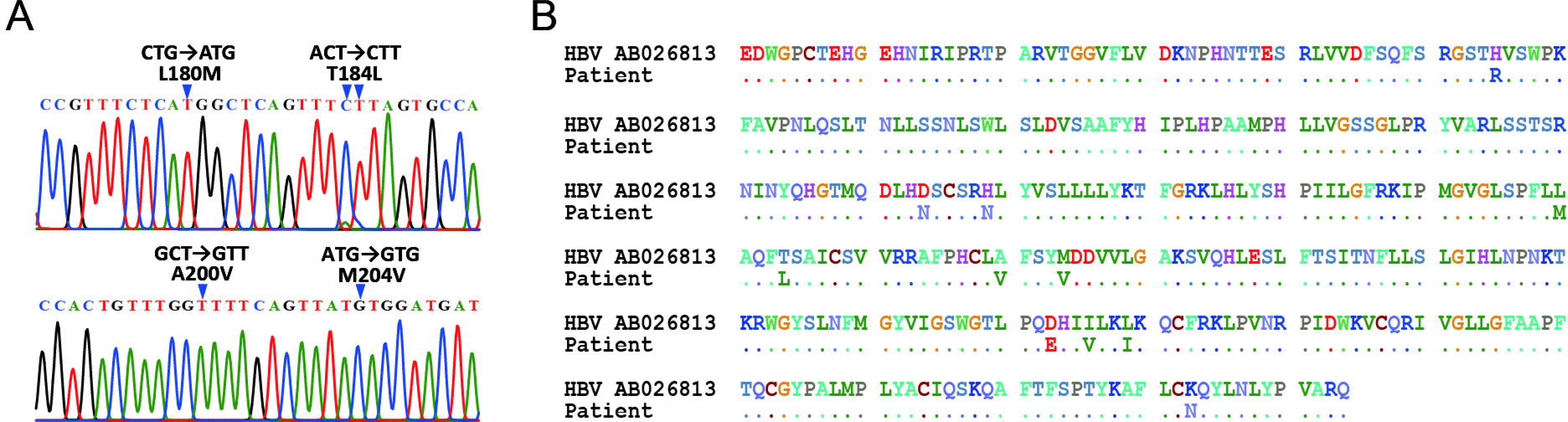
Mutations detected in the patient with HBV rebound. (A) Sequencing result of the sample collected in December 2016. Mutations and corresponding RT amino acids are indicated. (B) Amino acid changes in HBV in the patient compared with wild-type HBV strain AB026813.

Of note, as shown in Fig. 2A, the rtL180M/T184L/A200V/M204V quadruple mutant was a major viral population at breakthrough. Clonal sequencing further confirmed this result. Among 37 sequenced clones, 26 (70.3%) were rtL180M/T184L/A200V/M204V quadruple mutants. The other nine clones included two (5.4%) with rtL180M/M204V, three (8.1%) with rtL180M/T184L/M204V, five (13.5%) with L180M/A200V/M204V and one (2.7%) with T184L/A200V/M204V.

### Viral replication competence and TDF sensitivity of rtA200V containing mutants

To further investigate TDF sensitivity of this quadruple mutant, we generated HBV constructs containing HBV mutants including rtL180M/M204V, rtL180M/T184L/M204V, rtA200V and rtL180M/A200V/T184L/M204V. These constructs were transiently transfected into HepG2 cells and HBV replicative intermediates were analyzed. In agreement with results from previous studies, both rtL180M/T184L and rtL180M/T184L/M204V mutants showed lower replication competence (49.1% and 26.7%, respectively) than did the wild-type controls (Fig 3A and 3B). The replication competence of the rtA200V mutant and the rtL180M/A200V/T184L/M204V quadruple mutant was also attenuated (55.0% and 22.5% that of the wild-type, respectively; Fig 3A and 3B). Because these mutations were not located in the promoters of the HBV S and preC genes, no significant differences in the production of e and S antigens were observed (data not shown).

**Fig. 3.**
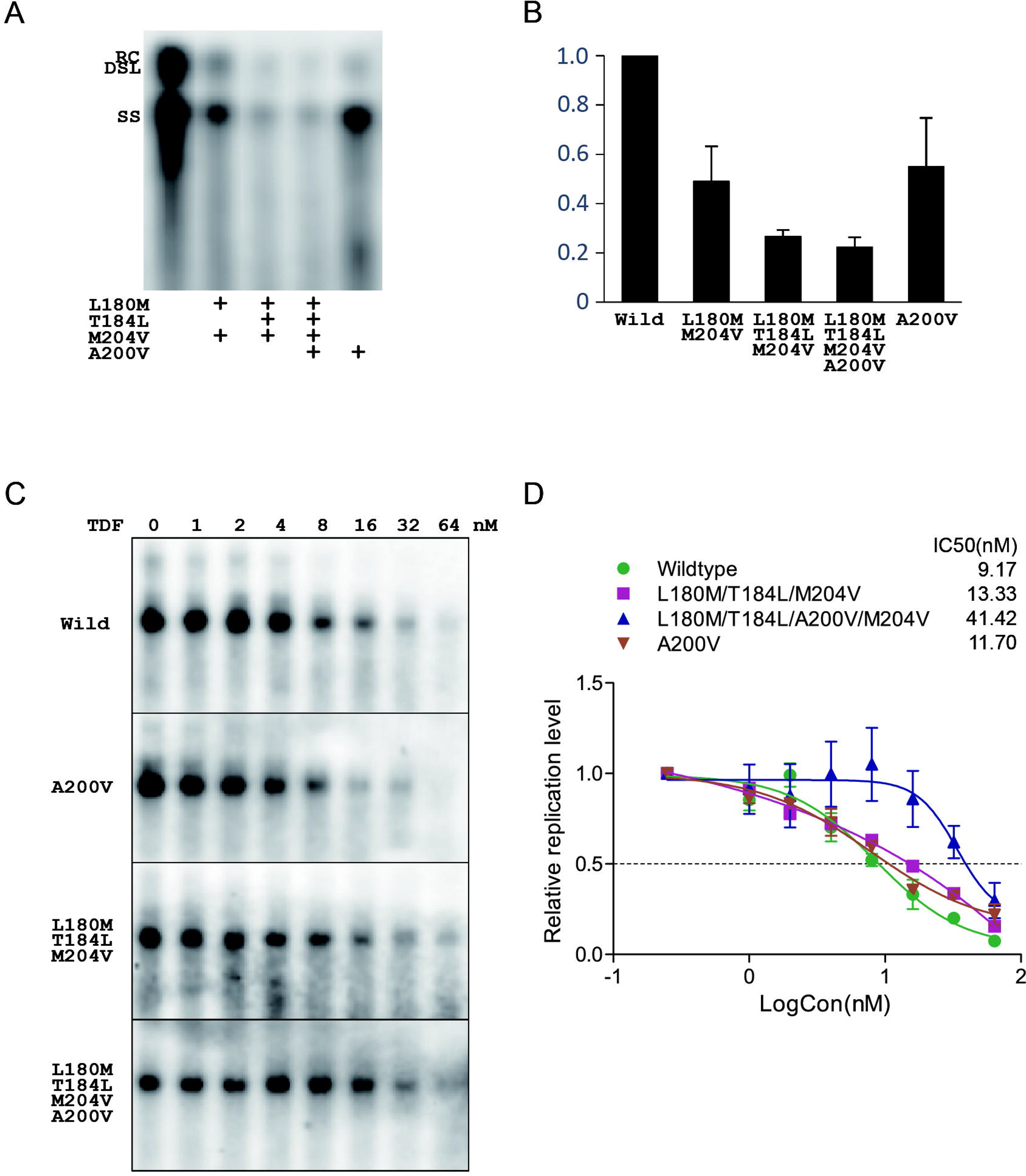
Virological profile of the TDF-resistant mutant. (A) Southern blotting of HBV mutants. RC, DSL, and SS DNA are indicated. (B) Quantitative analysis of the replication competence of the HBV mutants. HBV relative replication was determined through Southern blotting and analysis in Image Lab software. (C) Southern blotting of different mutants treated with different concentrations of TDF. (D) Dose-effect curves of different mutants. EC50 for TDF in different mutants are shown.

We treated the transfected HepG2 cells with a series of concentrations of TDF. HBV replication intermediate signals were collected, and the EC50 for TDF in each HBV mutant was determined by nonlinear regression with variable slope (Fig. 3C and 3D). In agreement with findings from previous reports, the rtL180M/T184L/M204V mutant was sensitive to TDF, with a 1.45-fold higher EC50 than that in the wild-type control (13.33 versus 9.17 nM), whereas the rtA200V mutant also displayed an EC50 comparable to that in wild type (1.28-fold, 11.70 nM). The EC50 of the rtL180M/T184L/A200V/M204V mutant was approximately 4.52 fold (41.42 nM) that in the wild-type control (Fig. 3D). Therefore, the quadruple mutant rather than the rtL180M/T184L/M204V ETV-resistance mutation or rtA200V single mutation led to moderate resistance to TDF.

To further specify the mutations conferring decreased susceptibility to TDF, we constructed six additional combinations of mutants: rtL180M/A200V, rtT184L/A200V, rtA200V/M204V, rtL180M/T184L/A200V, rtT180/A200V/M204V and rtT184L/A200V/M204V. Southern blot assays revealed that all constructs displayed decreased intracellular HBV replicative intermediates (Fig S1). All mutants expect rtT184L/rtA200V/rtM204V were sensitive to TDF, and had comparable EC50 values for TDF (Fig. S2). EC50 of rtT184L/rtA200V/rtM204V can not be determined because of the mutant’s extremely low replication competence. The above results further indicated that the rtL180M/A200V/T184L/M204V quadruple mutant conferred decreased susceptibility to TDF.

### Sensitivity of the quadruple mutant to ETV/TDF combination treatment

Because the clinical data indicated that the quadruple mutant was inhibited by ETV/TDF combination therapy, we next tested whether the quadruple mutant might be sensitive to ETV/TDF combination treatment *in vitro*. To this end, ETV and TDF were added into the medium at the EC50 (7.10 nM for ETV and 9.17 nM for TDF in our system) and with a 1.5-fold serial dilution. Both the wild type and the quadruple mutant were sensitive to ETV/TDF combination treatment (Fig. 4), even at the low concentration of 0.3 fold of each EC50.

**Fig. 4.**
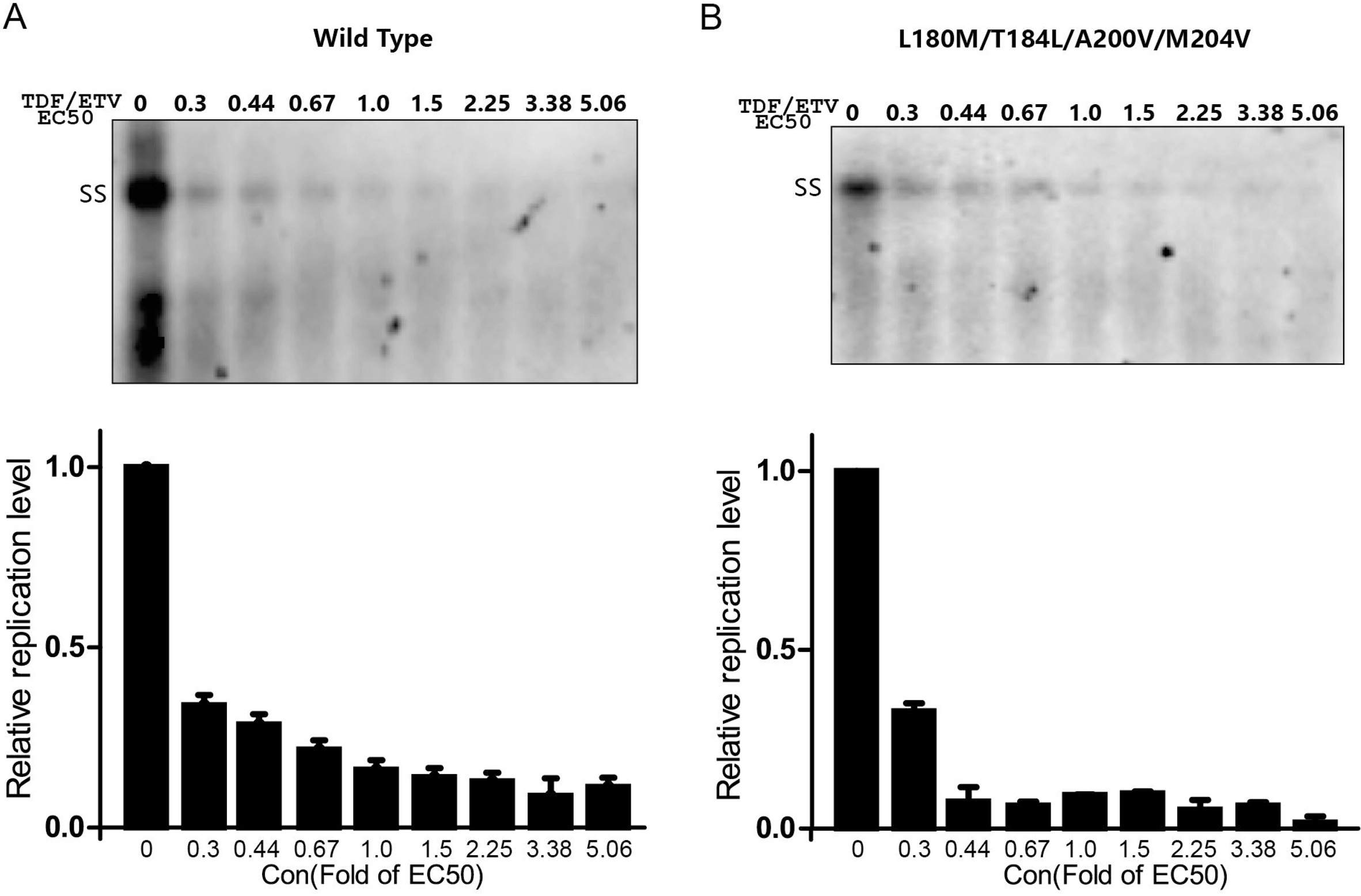
Sensitivity of the quadruple mutant to ETV/TDF combination therapy. Southern blotting and quantitative analysis of wild type HBV (A) and HBV quadruple mutants (B). ETV and TDF were added to the medium at the EC50 (7.10 nM for ETV and 9.17 nM for TDF in our system) and with 1.5-fold serial dilution.

### Spatial analysis of the quadruple mutant rtL180M/A200V/T184L/M204V

To illustrate the spatial position of the mutated rtA200V in HBV RT, we constructed a structural model of the HBV RT-DNA-tenofovir triphosphate complex with the PyMOL program. Previous studies have demonstrated that rtL180 and rtM204 are located near the catalytic and substrate binding center of RT, and their mutations affect the binding of substrates; rtT184 stabilizes the tyrosine-methionine-aspartate-aspartate (YMDD) loop, whereas rtT184L destabilizes the YMDD loop and thus hinders the binding of entecavir triphosphate (Fig. S3). rtA200 is located in the middle of the RT C domain loop, and is far from the YMDD motif, thus indicating that rtA200V might not directly affect binding and catalysis of substrates.

## Discussion

TDF based treatment results in high viral suppression in both NA-naïve patients and patients who have failed ETV, LAM/ADV and ETV/ADV therapy LAM-, ADV- or ETV-resistant mutants.(8–11, 15–18) More importantly, TDF also imposes a high genetic barrier to drug resistance. Few conserved mutations that are associated with TDF resistance have been observed during long-term clinical follow-up.(12) Sporadic virological breakthrough has been attributed to either poor patient compliance during treatment or episodes of late response.(10, 12) Several reported TDF-resistance mutations have failed to be confirmed.(19, 20) Recently, an rtS78T/sC69* mutant was reported to be TDF refractory but with only a slight increase in EC50.(13) Here, we identified a quadruple mutant of rtL180M/T184L/A200V/M204V in a patient who experienced LAM and ETV resistance and displayed viral rebound during TDF monotherapy. Moreover, our *in vitro* study demonstrated a 4.52-fold higher EC50 in this quadruple mutant compared with the wild-type virus.

Previous studies have shown that ETV resistance usually requires classical LAM-resistance mutations (rtL180M+rtM204V/I) together with additional mutations, such as rtT184, rtS202, rtI169 and rtM250.(5–7) Accordingly, LAM-resistant patients exhibit a markedly higher rate of ETV resistance than NA-naïve patients after 5-year treatment (51% vs.1.2%).(7) Similarly, our patient sequentially developed LAM resistance, ETV resistance and TDF resistance, and the TDF-resistance mutant harbored four mutations, three of which were classical ETV-resistance mutations (rtL180M/T184L/M204V). The possibility of TDF-resistance might be worthy of consideration in sequential treatment following the emergence of NA-resistance, especially ETV-resistance.

Our study also demonstrated the potency of TDF/ETV combination therapy for TDF-refractory patients. It was reported that the efficacy of TDF monotherapy is not inferior to that of TDF/ETV combination therapy in patients with MDR HBV.(21) In addition, withdrawal of NA in virologically suppressed MDR CHB patients undergoing TDF+NA (most ETV) combination therapy still has sustained anti-viral effects.(22) However, the viral breakthrough in our case occurred after a switch from ETV/TDF combination therapy to TDF monotherapy, indicating a potential risk of viral breakthrough after NA withdrawal. Fortunately, this patient with TDF resistance achieved virological suppression after TDF/ETV combination rescue therapy, thus suggesting that TDF/ETV combination therapy might be a promising choice for controlling potential TDF-resistant HBV. Because the nucleoside analogue ETV and nucleotide analogues TDF have different ribose structures, we speculated that they might function synergistically. This hypothesis was supported by the finding that both the quadruple mutant and the wild type virus were sensitive to *in vitro* ETV/TDF combination treatment. In agreement with our results, several clinical studies have also shown that TDF-refractory HBV replication can be gradually controlled by ETV/TDF combination therapy.(13, 23) In addition, we found that this quadruple mutant was sensitive to GLS4, a core assembling inhibitor (Jiang *et al*, unpublished data), thus highlighting the potential importance of developing other HBV inhibitors.

Lee *et al.* has reported rtA200V in one patient with NA sequential therapy.(23) However, rtA200V was not dominant and a number of other accompanying mutations were detected. Additionally, *in vitro* phenotypic analysis was not performed. rtA200V alone did not cause TDF-resistance, indicating that rtA200V might synergize with ETV-resistance mutations. rtL180 and rtM204 are known to be located near the catalytic and substrate binding center of RT, and their mutations affect substrate binding; rtT184 forms a hydrogen bond with the rtS202 side-chain, which sequentially forms hydrogen bonds with rtL180 and rtM204 to stabilize the YMDD loop.(24–26) Remodeling analysis indicated that rtA200 is located in the middle of the RT C domain loop, suggesting that rtA200V might not directly interfere with binding of tenofovir triphosphate. Instead, rtA200V might co-operate with rtT184L and further affect the spatial position of the YVDD loop, and subsequently impede binding of tenofovir triphosphate to the RT complex.

Our study was limited to a single case of a patient with TDF resistance. However, potential cases with TDF-resistant mutants might be underestimated because TDF-resistant mutants might be suppressed by ETV/TDF combination therapy. TDF-resistance associated mutation should be carefully investigated in patients with sequential NA treatment.

In summary, we report an rtL180M/T184S/A200V/M204V mutant that gained moderate resistance to TDF monotherapy in a stepwise manner. ETV/TDF combination therapy was found to be potently effective against this MDR HBV mutant.

## Abbreviations

TDF: tenofovir disoproxil fumarate
MDR: multidrug-resistant
HBV: hepatitis B virus
NA: Nucleoside and nucleotide analogs
LAM: lamivudine
ETV: entecavir
rt: HBV reverse transcriptase
EC50: half maximal effective concentration
ADV: adefovir dipivoxil
HBeAg: hepatitis B e antigen
HBsAg: hepatitis B surface antigen
CHB: chronic hepatitis B
QD: quaque die
ALT: alanine aminotransferase
cccDNA: covalently closed circular DNA
DMEM: Dulbecco’s modified Eagle’s medium
FBS: fetal bovine serum
ELISA: enzyme-linked immunosorbent assay
mRNA: messenger RNA
IU: international unit
RC: relaxed circular DNA
DSL: double-stranded linear DNA
SS: single stranded DNA
YMDD: tyrosine-methionine-aspartate-aspartate

## Conflict of interest

No potential conflicts of interest were disclosed

## Authors contributions

Hui Zeng and Xianbo Wang designed the study; Dong Jiang, Jianghua Wang, Xuesen Zhao and Chuan Song completed the lab experiment; Yuxin Li and Qun Zhang collected the clinical data; Dong Jiang and Jianghua Wang processed the data and prepared the draft of manuscript; Dong Jiang, Jianghua Wang, Xuesen Zhao, Hui Zeng and Xianbo Wang extensively discussed and finalized the manuscript.

## Acknowledgment

The authors thank Dr. Hong You and Dr. Hong Zheng for scientific guidance and discussion. The authors thank Dr. Jie Guo for clinical data collection.

## Notes

Funding: Supported by Beijing Municipal Administration of Hospitals Clinical Medicine Development of Special Funding Support (ZYLX201707) and the Natural Science Foundation of China (81572043).

