## Supplemental Figure Legends for "Entecavir resistance mutations rtL180M/T184L/M204V combined with rtA200V lead to tenofovir resistance"

**Fig. S1. Replication competence of A200V containing mutants.** (A) Southern blotting of HBV mutants. SS DNA are indicated. (B) Quantitative analysis of the replication competence of different HBV mutants. HBV relative replication was determined by Southern blotting and analysis in Image Lab software.

**Fig. S2. Susceptibility of different A200V containing mutants to TDF treatment.** (A) Southern blotting of different mutants treated with different concentrations of TDF. (B) Dose-effect curve of different A200V containing mutants. EC50 values for TDF in different mutants are shown.

**Fig. S3. Molecular modeling of HBV RT.** A model of the HBV polymerase-DNA-tenofovir triphosphate complex was constructed. Positions corresponding to rtL180M/T184L/A200V/M204V are indicated.

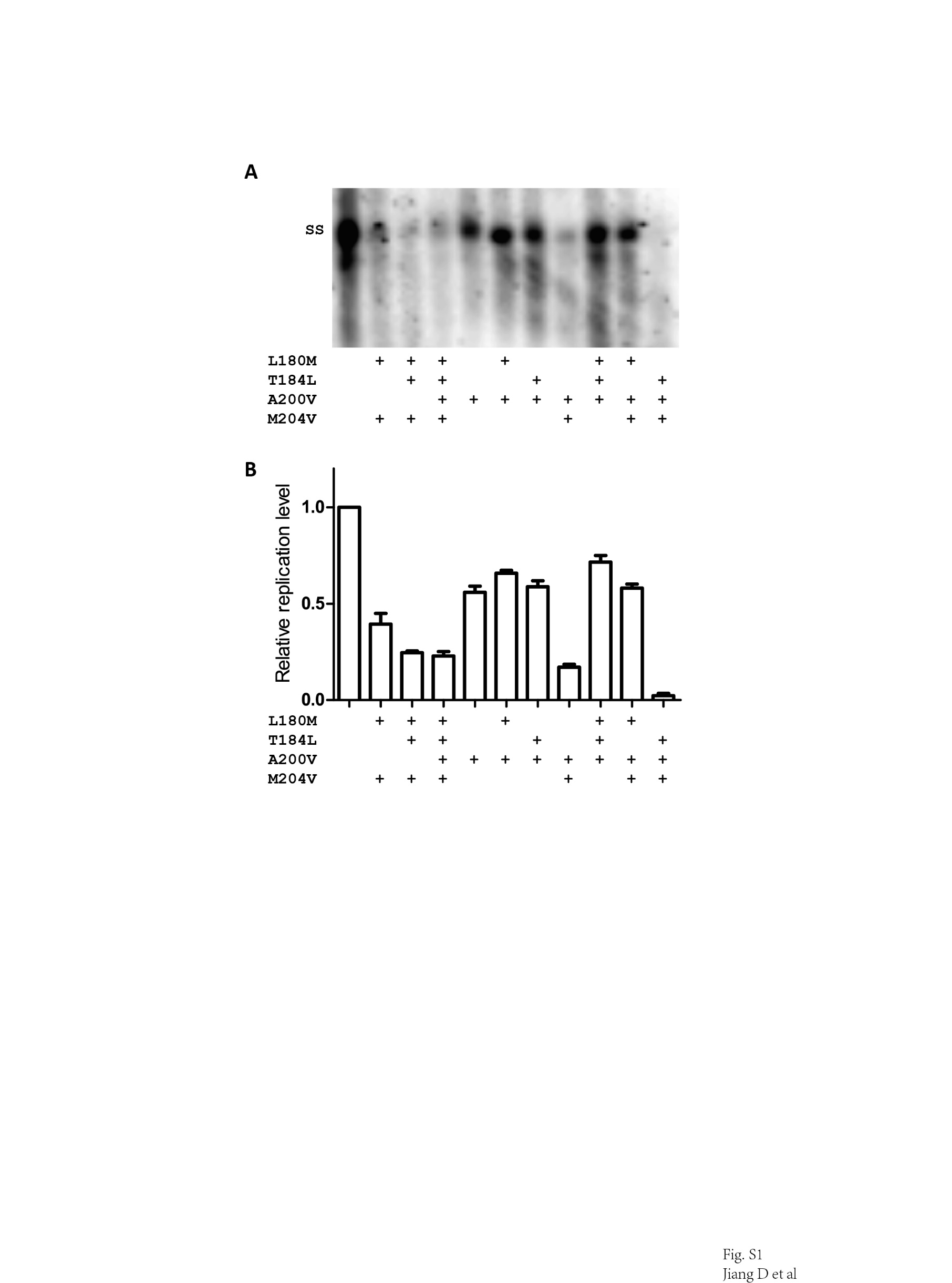

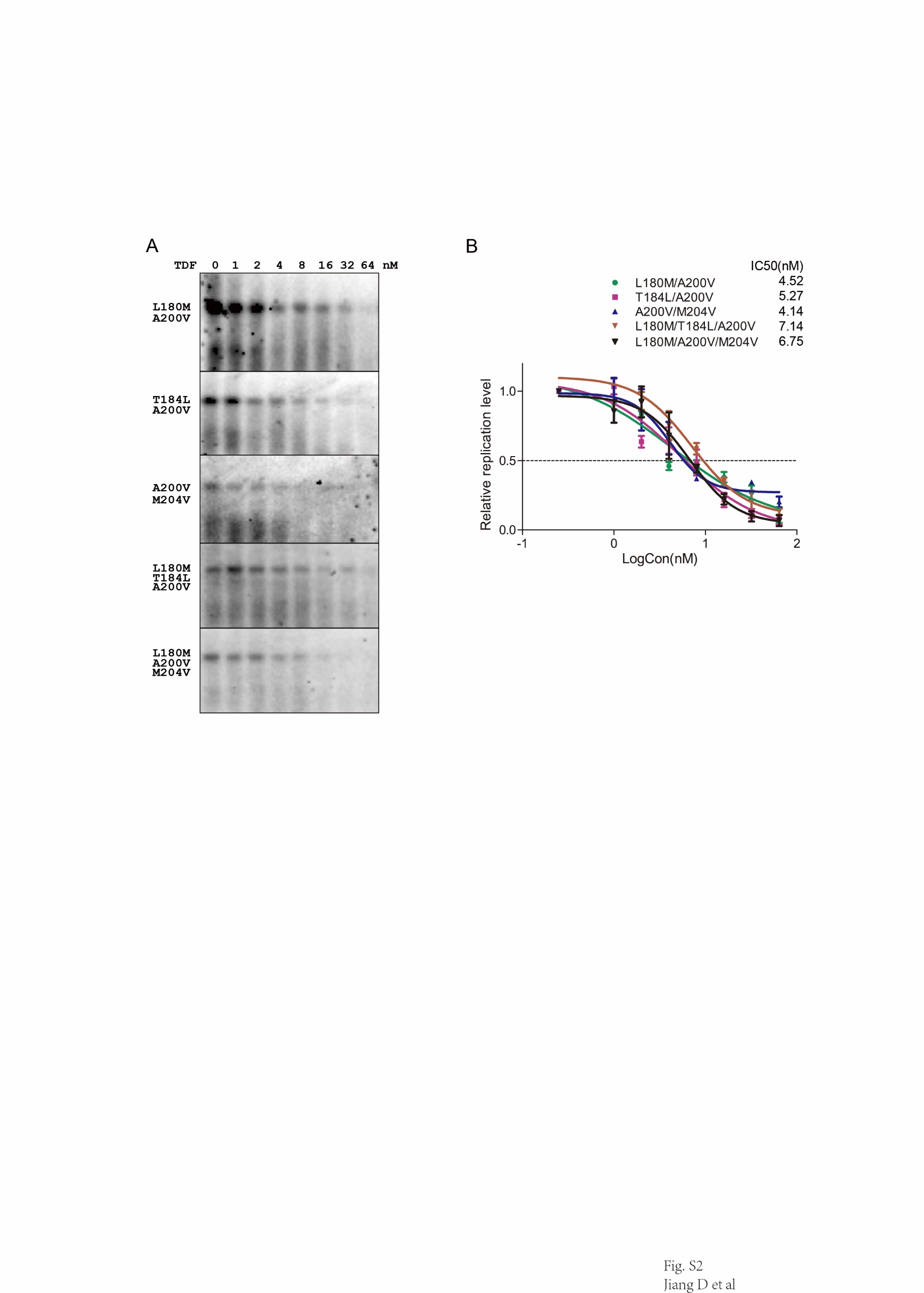

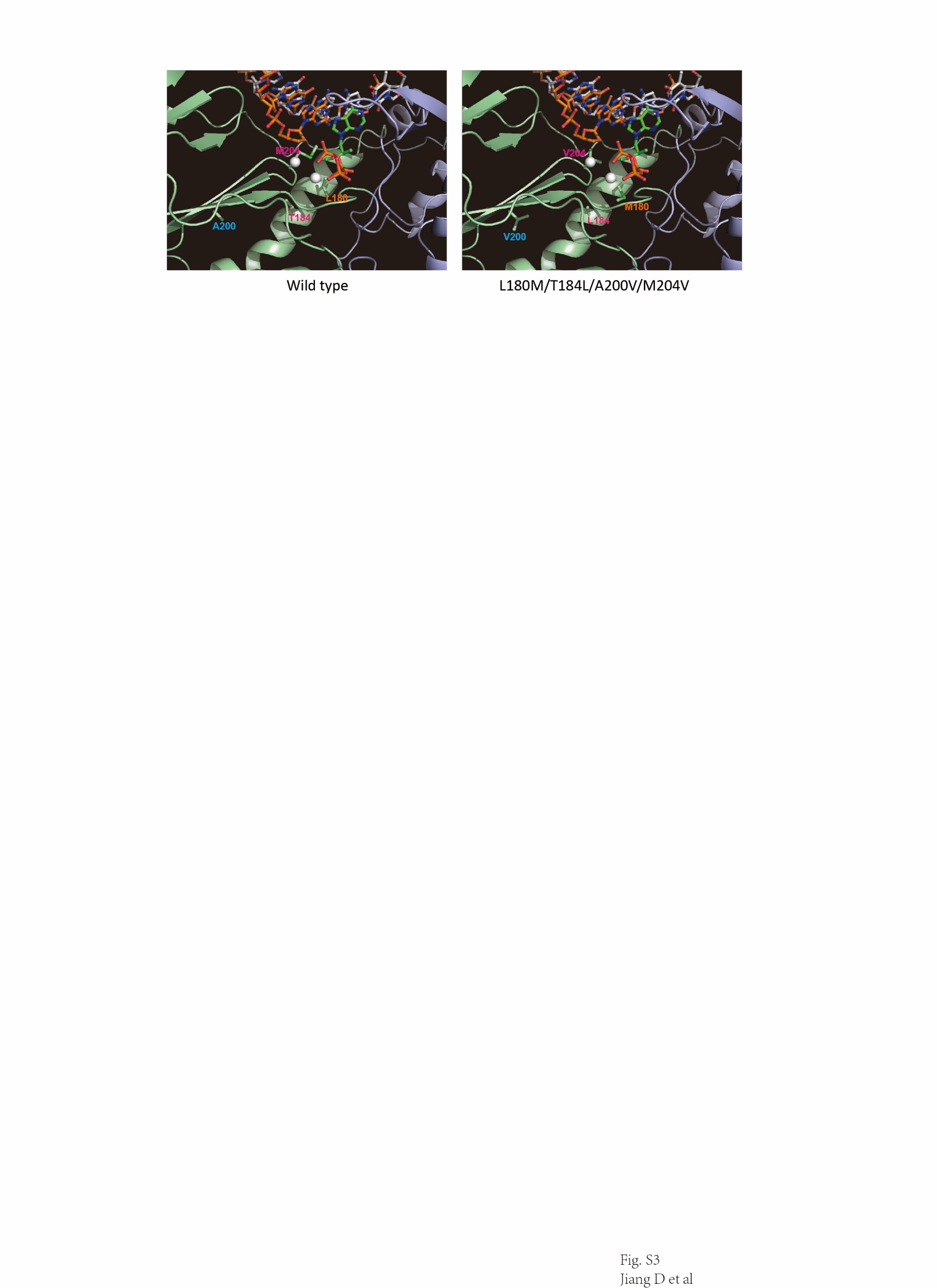
