## Supplementary figures and images for "Entecavir resistance mutations rtL180M/T184L/M204V combined with rtA200V lead to tenofovir resistance"

### Fig. S1

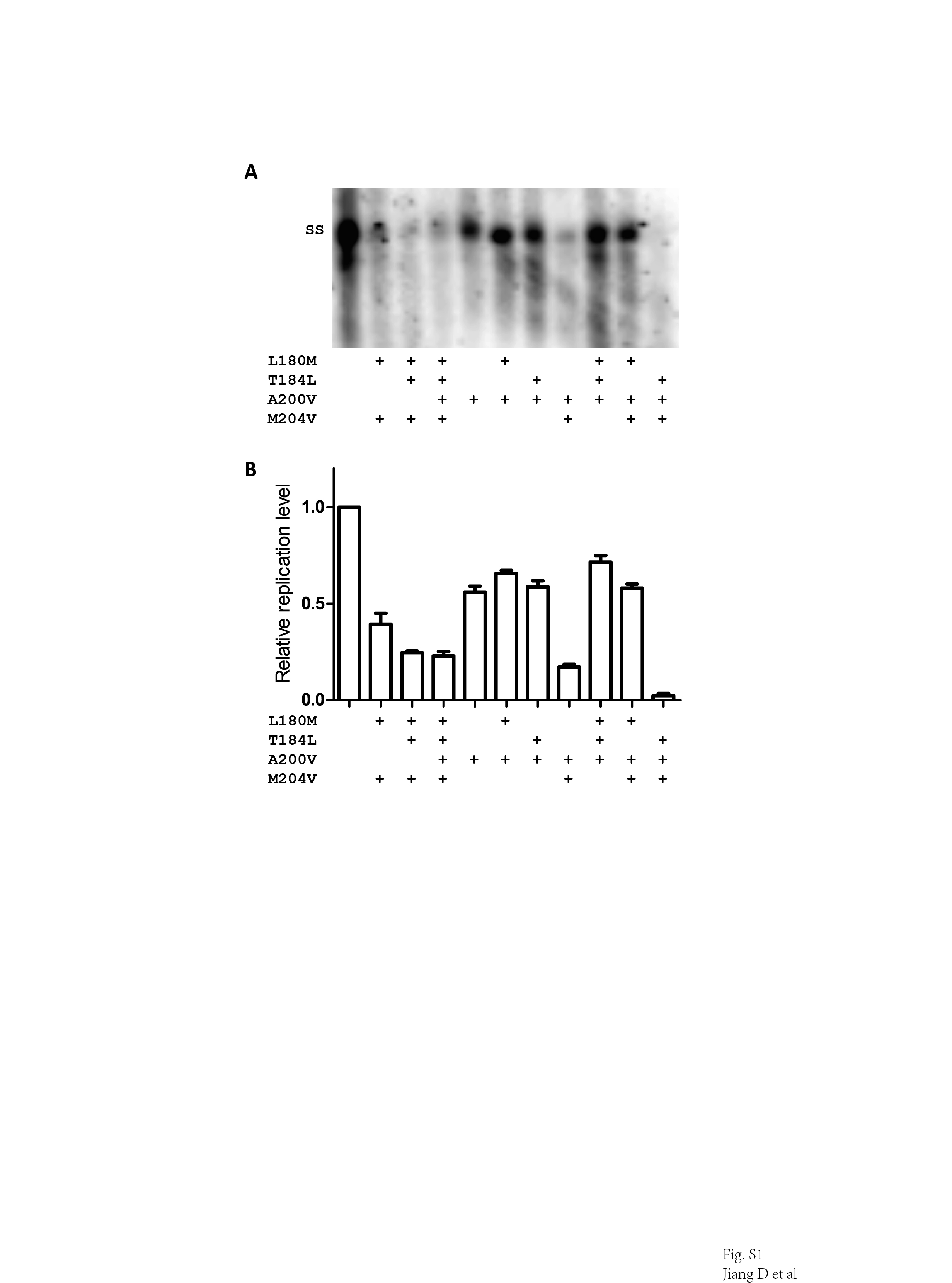

### Fig. S2

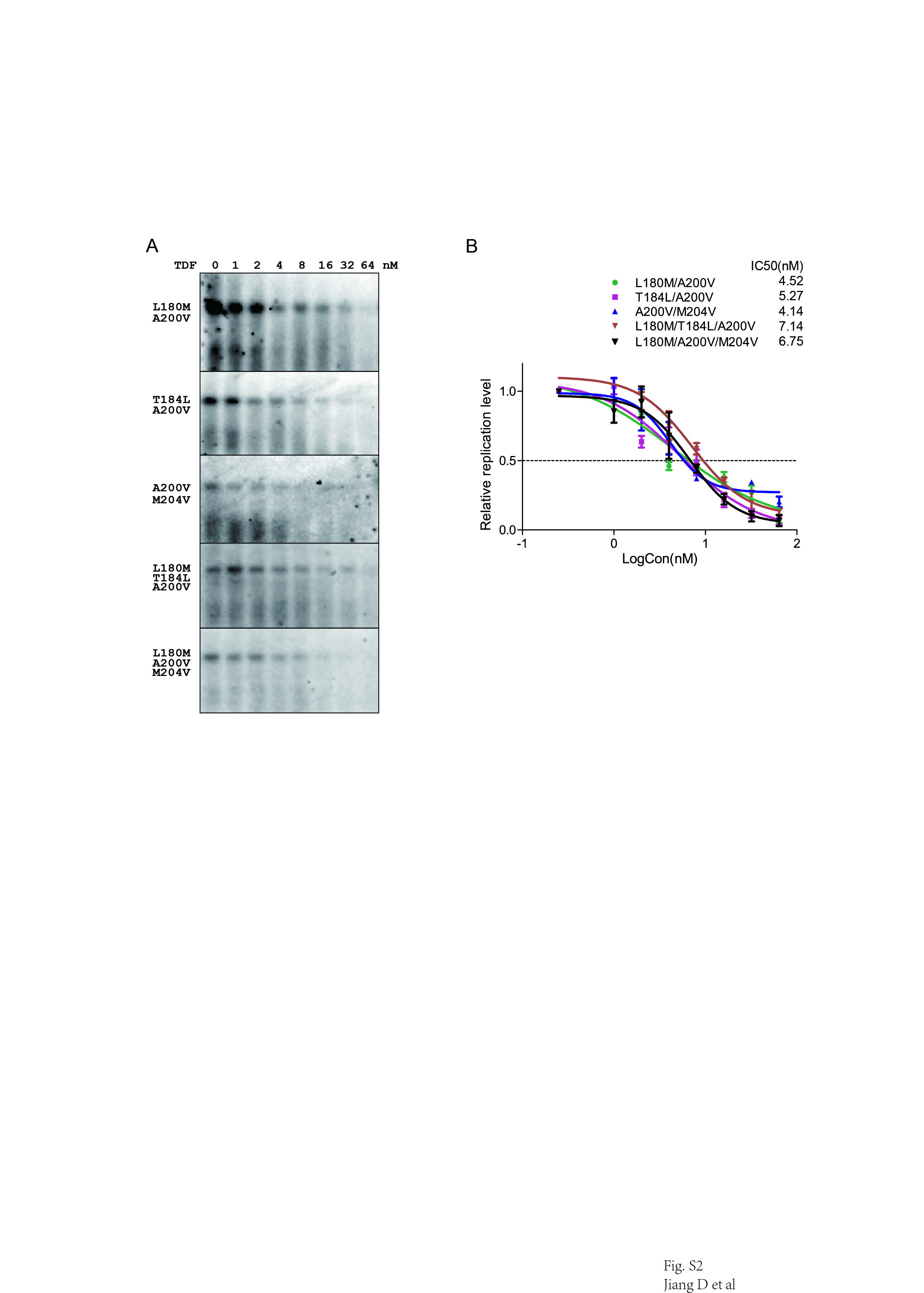

### Fig. S3

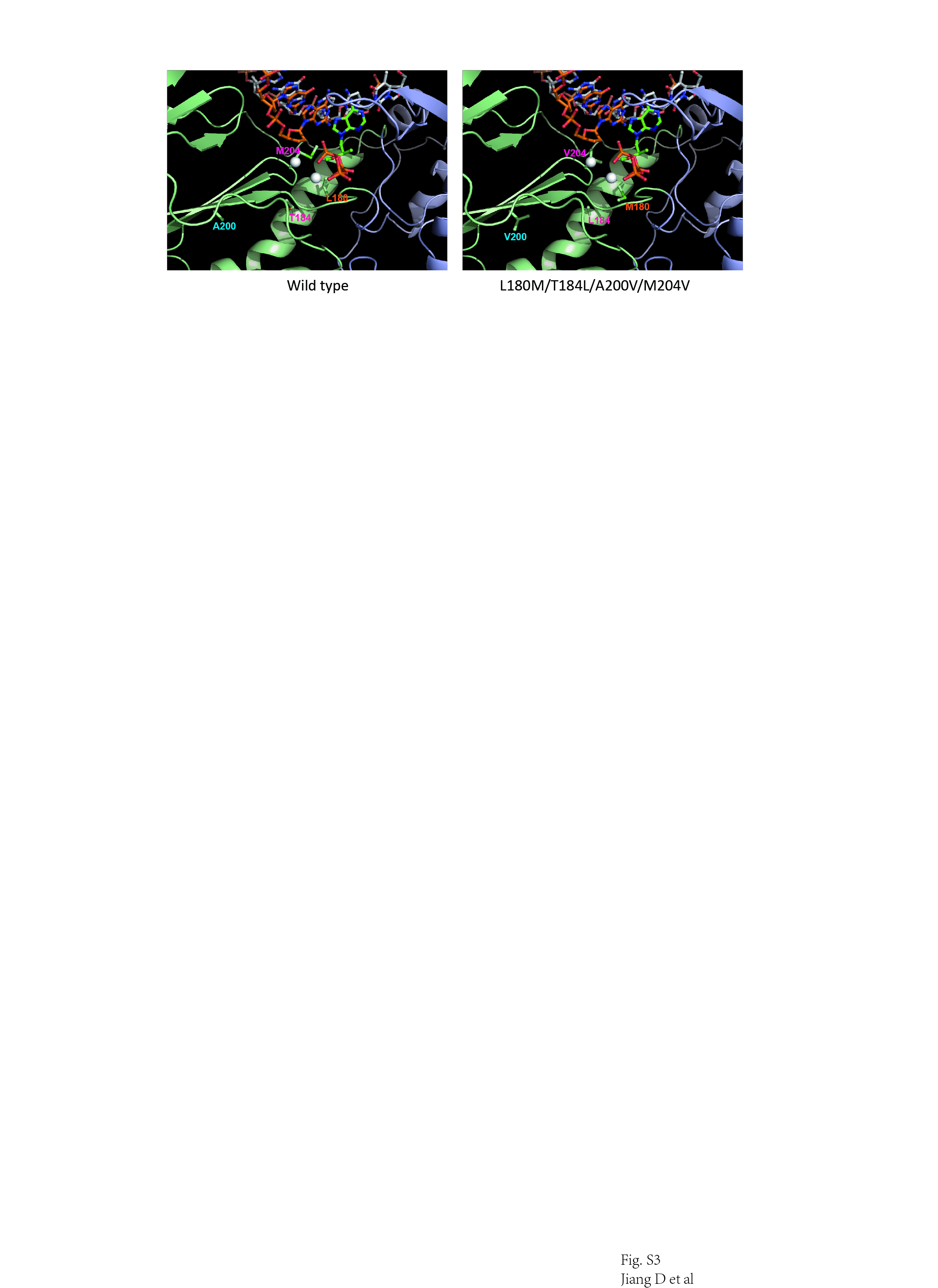
